# Andexanet alfa-associated heparin resistance in cardiac surgery: mechanism and *in vitro* perspectives

**DOI:** 10.1101/2024.09.09.612152

**Authors:** Charlene V. Chabata, Haixiang Yu, Lei Ke, James W. Frederiksen, Prakash A. Patel, Bruce A. Sullenger, Nabil K. Thalji

## Abstract

**Background:** Andexanet alfa (andexanet) is the only FDA-approved antidote for direct factor Xa (FXa) inhibitors but has been reported to cause resistance to unfractionated heparin (UFH). This has delayed anticoagulation for procedures requiring cardiopulmonary bypass (CPB). The mechanism, andexanet and UFH dose dependence, and thrombotic risk of andexanet-associated heparin resistance are unknown.

**Methods:** The effect of andexanet *in vitro* was determined using activated clotting times (ACT) and thromboelastography (TEG). *Ex vivo* CPB circuits were used to determine whether andexanet impaired anticoagulation for extracorporeal circulation. Kinetics of antithrombin (AT) inhibition of FXa and thrombin were measured in the presence of andexanet. Equilibrium modeling and thrombin generation assay (TGA) validation were used to predict the role of andexanet, AT, and UFH concentrations in andexanet-associated heparin resistance.

**Results:** Andexanet prevented UFH-mediated prolongation of ACT and TEG times. At lower concentrations of andexanet, heparin resistance could be overcome with suprapharmacologic doses of UFH, but not at higher andexanet concentrations. Andexanet rendered standard doses of UFH inadequate to prevent circuit thrombosis, and suprapharmacologic UFH doses were only partially able to overcome this. Scanning electron microscopy demonstrated coagulation activation in circuits. Andexanet prevented UFH enhancement of AT-mediated inhibition of FXa and thrombin. Equilibrium modeling and TGA validation demonstrated that andexanet creates a triphasic equilibrium with UFH and AT: initial UFH unresponsiveness, normal UFH responsiveness when andexanet is depleted, and finally AT depletion. Sufficient CPB heparinization can only occur at low therapeutic andexanet doses and normal AT levels. Higher andexanet doses or AT deficiency may require both AT supplementation and very high UFH doses.

**Conclusions:** Andexanet causes heparin resistance due to redistribution of UFH-bound AT. If andexanet cannot be avoided prior to heparinization and direct thrombin inhibitors are undesirable, our *in vitro* study suggests excess UFH should be considered as a potential strategy prior to AT supplementation.

**Highlights:**

- Andexanet alfa causes heparin resistance not by depleting antithrombin, but rather by sequestering heparin-bound antithrombin such that it cannot act as an anticoagulant.
- Heparin responsiveness in the presence of Andexanet alfa is triphasic such that the effect of a dose of heparin can now be predicted in vitro based on the relative concentrations of andexanet, heparin, and antithrombin.
- The in vitro insights provided by this work provide a rational starting point for further clinical elucidation of the problem and management of andexanet-associated heparin resistance

## Introduction

Direct factor Xa (FXa) inhibitors are widely prescribed oral anticoagulants for prevention and treatment of venous thromboembolism and prevention of stroke in patients with atrial fibrillation.^1–6^ Like other anticoagulants, direct FXa inhibitors can cause bleeding.^7, 8^ In 2018, Andexanet alfa (andexanet) became the only FDA-approved reversal agent for direct FXa inhibitors.^9^ Andexanet is a recombinant variant of FXa that is catalytically inactive due to mutation of the active site serine to an alanine (S195A, chymotrypsin numbering system^10^).^11–15^ Andexanet also lacks the membrane-binding γ-glutamic acid (gla) domain which would otherwise cause andexanet to interfere with assembly of the prothrombinase complex. Because andexanet retains the active site structure of FXa, it functions as a drug-sequestering antidote to direct FXa inhibitors by binding tightly to direct FXa inhibitors.^11^ However, despite this tight binding, andexanet is administered at high doses (micromolar plasma concentrations) because its mechanism of action requires 1:1 stoichiometry with the anticoagulant.^12–16^

Cardiopulmonary bypass (CPB) poses a unique hemostatic challenge. Exposure of blood to extracorporeal membranes results in contact activation, surgical tissue injury sweeps coagulation activators into the bypass circuit, and a significant portion of a patient’s blood remains static in a large cardiotomy reservoir.^17–20^ For this reason, anticoagulation with unfractionated heparin (UFH) is required to prevent thrombosis. UFH administration usually results in a predictable degree of anticoagulation, but in some instances, collectively described as “heparin resistance,” the level of anticoagulation achieved from a dose of UFH is lower than that dose is expected to produce. This has been attributed to a variety of causes, including high factor VIII levels, and nonspecific plasma protein binding. Perhaps the best-known cause of heparin resistance, however, is antithrombin (AT) deficiency. UFH exerts its anticoagulant effects by potentiating the activity of AT.^21^ In the absence of heparin, AT rapidly and irreversibly inhibits thrombin, FXa, and FIXa, and is a key regulator of coagulation. UFH increases the rate of AT-mediated inactivation of its targets by up to 4000-fold. Since AT is necessary to achieve a heparin anticoagulant effect, AT deficiency has been associated with heparin resistance.^22^ While the precise definition of heparin resistance in cardiac surgery is subject to much debate, the International Society on Thrombosis and Haemostasis (ISTH) scientific subcommittee on perioperative and critical care thrombosis and hemostasis recently proposed the definition of the inability of 500 U/kg to produce an activated clotting time (ACT, a common point-of-care coagulation test) of at least 480 seconds.^23, 24^ Heparin resistance due to AT deficiency is usually managed with AT supplementation, as there is typically sufficient heparin present but insufficient AT.

Since approval of andexanet, several case reports, mostly in cardiac and vascular surgery, have described a potentially new form of heparin resistance following andexanet administration.^25–31^ In some cases, achieving therapeutic anticoagulation for CPB or other procedures was possible but required suprapharmacologic doses of UFH in excess of 500 U/kg.^31^ In others, high doses of UFH were not sufficient, suggesting that there are multiple factors that determine heparin responsiveness following administration of andexanet.^27–30^ Since rapid therapeutic anticoagulation is critical in the peri-procedural setting, identification of the factors that affect heparin responsiveness is crucial to determining the optimal strategy for post andexanet anticoagulation. This is a particularly pressing issue, since current STS/SCA/AmSECT/SABM guidelines recommend administration of andexanet prior to emergent cardiac surgery in patients with recent rivaroxaban or apixaban administration, with no mention of andexanet-associated heparin resistance.^32–34^

It is unknown if andexanet-associated heparin resistance occurs via the same mechanism as AT-mediated heparin resistance. Indeed, case reports of overcoming andexanet-associated heparin resistance with very high doses of UFH indicates that some patients do, in fact, have sufficient AT to allow for an anticoagulant effect.^31^ Early reports of andexanet described it as having the ability to reverse the effects of heparinoids due to its ability to bind AT in their presence.^11^ Thus, we hypothesize that andexanet-associated heparin resistance is not due to true AT deficiency, but rather redistribution of AT that renders it ineffective as an anticoagulant.

Given the reports of andexanet-associated heparin resistance, understanding the nature of the problem is crucial. Here, we characterize andexanet-associated heparin resistance with clinical coagulation tests and extracorporeal circulatory models. We show that andexanet-associated heparin resistance is distinct from heparin resistance due to AT deficiency and enumerate the factors that govern heparin responsiveness after andexanet administration.

## Methods

All data and supporting materials have been provided with the published article.

### Reagents and recombinant proteins

Reagent sourcing and recombinant protein production is summarized in the Supplemental Methods.

### Human whole blood samples

For human whole blood studies performed at the Children’s Hospital of Philadelphia (activated clotting times and antithrombin level measurements), the institutional review board (IRB) of the Children’s Hospital of Philadelphia Research Institute approved the human phlebotomy and blood use performed. Informed consent was obtained from all subjects. For these studies, 10 mL whole blood was collected from 8 healthy human donors in one-tenth volume of 3.2% sodium citrate.

For human whole blood studies performed at Duke University, informed consent was obtained from each healthy human adult volunteer under a Duke University IRB approved protocol. All procedures were in accordance with institutional guidelines.

### Activated Clotting Time (ACT) Measurements

45 μL citrated human whole blood was mixed with UFH, andexanet, and rivaroxaban to the specified concentrations. Samples were recalcified to 10 mM CaCl_2_. The total blood volume following additions was 50 μL with 20 mM HEPES, 150 mM NaCl_2_, pH 7.4. ACT was then measured using a Hemochron Signature Elite ACT analyzer with Hemochron Jr. ACT+ cuvettes per the manufacturer’s instructions. For ACT measurements from *ex vivo* circuit blood, blood from the circuit was periodically sampled and analyzed as above.

### Thromboelastography (TEG)

320 μL of citrated whole blood from healthy volunteers was mixed with UFH and rivaroxaban (total volume 10 μL in 20 mM HEPES, 150 mM NaCl, 2 mM CaCl_2_). 10 μL kaolin solution and 10 μL buffer or andexanet at varying concentrations was added. The reaction was initiated with 10 mM CaCl_2_ in a TEG cup and clot elasticity was monitored at 37 °C in a Haemonetics Thromboelastograph Analyzer according manufacturer’s instructions.

### Determination of the kinetics of FXa-AT complex formation

Andexanet and UFH were added to recalcified (5 mM CaCl_2_ final) human FX-deficient plasma at room temperature along with 1.33 mM GPRP and 1 μM dabigatran to prevent clotting. 25 nM FXa was added to aliquots of the plasma mixture at different time points in a reverse time course. Samples were quenched simultaneously with 50 μM B-EGRCK and 50 μg/mL polybrene. incubated for 10 minutes, and then diluted tenfold with PBS, 0.1% Tween-20, 6% BSA, pH 7.4. FXa-AT standards (0-30 nM) were prepared as previously described.^35^ FXa-AT complex levels were then measured using a previously described ELISA.^35^ Samples were analyzed with a SpectraMax 190 microplate reader (Molecular Devices) and FXa-AT concentrations were determined using the generated standard curve.

### Determination of the kinetics of TAT complex formation

Andexanet was added to a mixture of 3.1 μM AT, 4 U/mL UFH, and 5 mM CaCl_2_ in 20 mM HEPES, 150 mM NaCl, pH 7.4. 500 pM plasma-derived human thrombin was then added to aliquots of the mixture at different time points in a reverse time course, and all samples were quenched simultaneously with 50 μM FGRCK (PPACK). Samples were then analyzed for TAT content with the Enzygnost TAT micro kit according to the manufacturer’s instructions.

### Equilibrium modeling of the interaction between Andexanet, UFH, and AT

Equilibrium concentration of UFH-AT complexes was modeled using KinTeK Explorer (KinTeK Corp.). The model was run with varying UFH concentrations in 0.25 μM increments from 0 to 7 μM UFH, and the equilibrium UFH-AT concentration (taken as the UFH-AT concentration at 60 seconds) was plotted versus UFH concentration. The model was run subject to the following expressions:

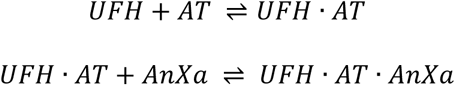

The second order UFH/AT association rate constant (1.7 x 10^7^ M^-1^s^-1^) and the first order UFH-AT dissociation rate constant (0.2 s^-1^) were used for the first expression.^36^ The second order UFH-AT/Andexanet association rate constant (8.2 x 10^4^ M^-1^s^-1^) and the first order UFH-AT- andexanet dissociation rate constant (0.05 s^-1^) were used for the second expression.^37^ The rate constants used for the second expression were from a study of gla-domainless FXa^S195A^ (biochemically identical to andexanet) binding to the AT-heparin pentasaccharide complex.^11, 35^

### Thrombin generation assays (TGAs)

TGAs were performed in normal human plasma as previously described^38^ with a modification to accommodate the addition of UFH, andexanet, and supplemental AT, as well as a modification to decrease the sensitivity to UFH (see Detailed Methods).

### Ex Vivo Membrane Oxygenator Circuit

The *ex vivo* membrane oxygenator circuit used in this study has been described before, and is summarized in the Supplemental Methods.^20, 39^ After setup, the circuit was primed with phosphate buffered saline (PBS) and circulated at 50 mL/min for ∼30mins. In this time, ∼50 mL of blood was drawn from a healthy donor. Immediately after, 80 μL of un-anticoagulated blood was used for assessment of baseline ACT. 4.5 mL of blood was anticoagulated with 3.2% sodium citrate while the rest was anticoagulated with 4 U/mL UFH or UFH plus rivaroxaban. Thereafter 5 mL was set aside for subsequent analyses and the 40 mL was used in the circulation experiments.

Once the PBS was drained from the circuit, the remaining 40 mL of heparinized blood was continuously circulated in the oxygenator circuit for 180 minutes or until the circuit thrombosed. 27 mL of this blood were used to initiate the procedure and the remaining 13 mL to replenish blood used for time-point aliquots. Andexanet was added after 60 minutes of circulation and monitored for an additional 120 minutes or until the circuit failed. During the study, samples were taken out for ACT assessment. Once the experiment was completed, the blood was collected for subsequent studies. Samples were anticoagulated with 3.2% sodium citrate and centrifuged at 3000 rpm for 10 minutes to give PPP. D-dimer levels were measured using the Asserachrom D-Di kit per the manufacturer’s instructions. Membrane samples were excised, rinsed in PBS, then prepared for and analyzed with SEM as described previously.^39^ *Quantification of AT levels in human plasma*

Pooled normal human plasma and healthy volunteer blood samples were analyzed for AT levels. For healthy volunteer samples, citrated blood was centrifuged at 190*g* for 15 minutes followed by a second centrifugation of the supernatant at 1,100*g* for another 15 minutes to produce platelet poor plasma. The plasma samples were then analyzed for AT levels using the Liatest ATIII kit per the manufacturer’s instructions.

### Statistical analysis

Experimental data were analyzed using Graphpad Prism 9. ACT and TEG measurements were analyzed with unpaired two-sample T-tests (**Fig. 1**). Factor Xa-AT formation and TGA individual data points are presented as mean±SEM as statistical comparisons between each data point do not provide relevant information.

**Figure 1.**
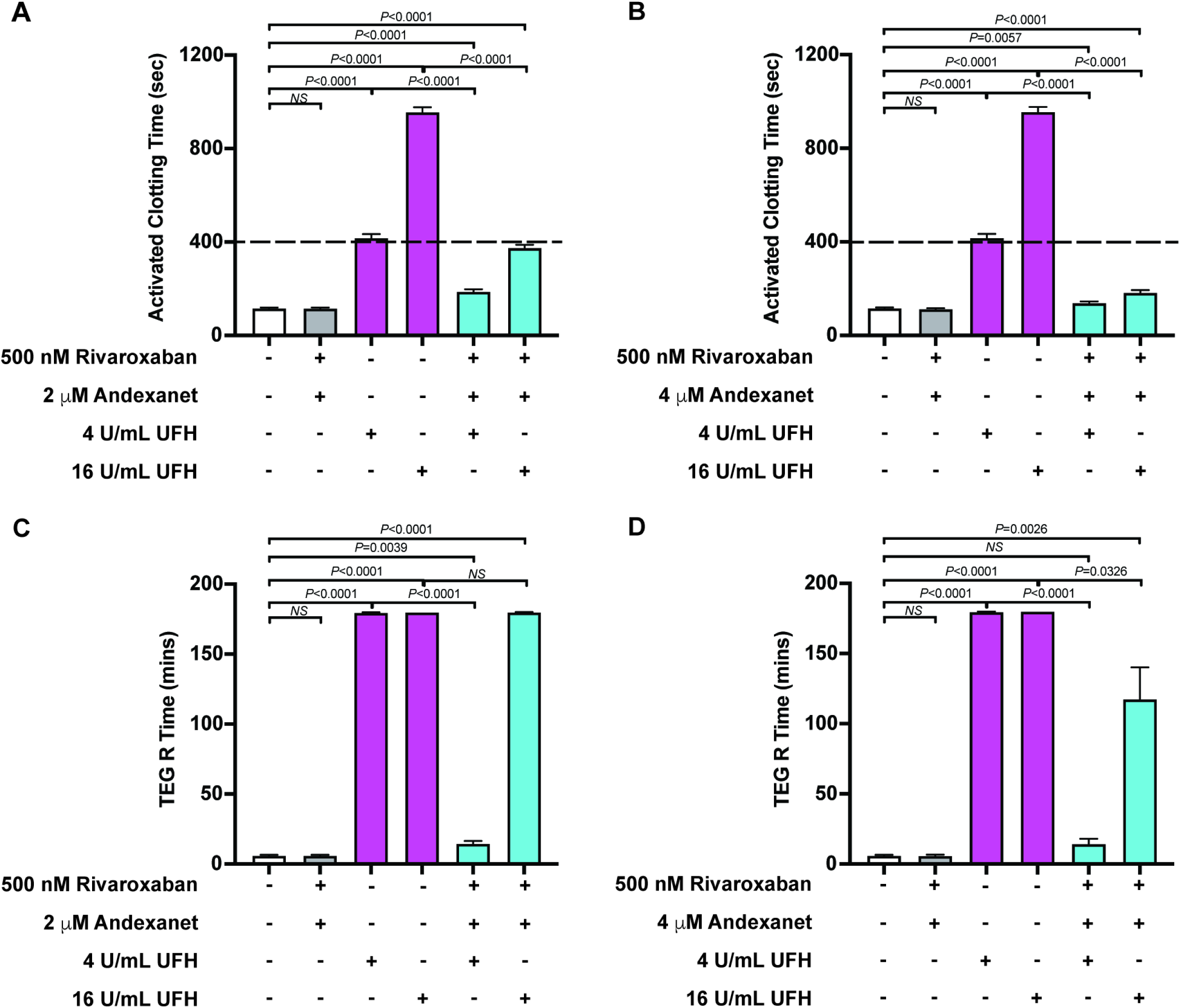
Effect of Andexanet on *in vitro* assessments of heparinization in whole blood. (***a, b***) ACT results following heparinization of human whole blood supplemented with (***a***) 2 μM andexanet (low-dose) or (***b***) 4 μM andexanet (high dose). The dashed line represents an ACT of 400 seconds, a commonly used ACT minimum for CPB. (***c,d***) TEG reaction (R) times following heparinization of human whole blood supplemented with (***c***) 2 μM andexanet or (***d***) 4 μM andexanet. TEG experiments were followed for 180 minutes, so any R times recorded as 180 minutes did not clot during the experiment. All data in panels ***a*** and ***b*** represent the mean ± SD of results from 8 separate donors, with each condition performed in duplicate. All data in panels ***c*** and ***d*** represent the mean ± SD of results from 4 separate donors.

## Results

### Andexanet prevents therapeutic prolongation of in vitro coagulation assays by UFH

Periprocedurally, the ACT is used to monitor the anticoagulant effects of UFH. To evaluate andexanet’s effects on UFH anticoagulation, we supplemented whole blood from healthy volunteers (n=8) with UFH and andexanet and measured ACTs. We also supplemented blood with 500 nM rivaroxaban which approximates peak rivaroxaban plasma concentrations.^40^ The donor whole blood demonstrated no overt coagulation abnormalities, and the mean AT concentration was 3.1 μM (**Supplementary Table S1**), identical to that measured in pooled normal human plasma. As expected, UFH at typical concentrations used for CPB (4 U/mL) prolonged the ACT beyond 400 sec, consistent with many institutions’ typical target ACT for CPB (**Fig. 1a, Supplementary Fig. S1a**).^41–43^ Supplementation of 2 μM andexanet (the expected plasma andexanet concentration after a 400 mg bolus with no infusion^16^) alone had no effect on the ACT, but in the presence of 4 U/mL UFH, andexanet markedly impaired UFH’s anticoagulant effects.

Heparin resistance in cardiac surgery can occur due to a number of factors. Regardless of the cause, initial treatment of subtherapeutic heparinization usually involves administration of additional UFH.^22^ Thus, to determine if high doses of UFH could overcome the heparin resistance, we repeated these experiments with 16 U/mL UFH. This resulted in ACT values that just approached the usual therapeutic target range, despite the fact that, in the absence of andexanet, 16 U/mL UFH severely prolongs the ACT (**Fig. 1a**). This suggests that heparin resistance due to low dose andexanet may be overcome with additional UFH administration, albeit with a very narrow therapeutic margin.

Andexanet is administered at one of two doses to match the reversal agent dose to the expected plasma concentration of the anticoagulant.^44^ To understand the effects of higher andexanet concentrations on heparin sensitivity, we measured ACTs in whole blood supplemented with 4 μM andexanet (the concentration of andexanet expected after an 800 mg bolus^16^). 4 U/mL UFH minimally prolonged the ACT (**Fig. 1b**). Unlike lower dose andexanet, in the presence of high dose andexanet, an adequate ACT could not be obtained even with high dose UFH (16 U/mL).

To ensure that our observations of andexanet-mediated heparin resistance were not assay-dependent, we assessed the effects of andexanet on thromboelastography (TEG)-monitored UFH anticoagulation. Contact-activated TEG is exquisitely sensitive to the effects of UFH, as demonstrated by the fact that doses at or above 1 U/mL UFH prolong the TEG R time beyond the limit of the test (>180 minutes, **Supplementary Fig. S2**). Consistent with this observation, UFH prolonged the TEG R time beyond the limit of the test (>180 minutes) at typical CPB and excess UFH doses (**Fig. 1c-d, Supplementary Fig. S1b**). In the presence of low dose andexanet, the TEG R time was only modestly prolonged by typical UFH doses. Based on the standard curve shown in **Supplementary Fig. S2**, this represents a level of anticoagulation less than what would be achieved with <0.25 U/mL UFH in the absence of andexanet. As in the ACT assay, excess UFH was seemingly able to overcome the andexanet-mediated heparin resistance (**Fig. 1c**), although the true level of anticoagulation could not be ascertained due to the high sensitivity of the assay. However, high dose andexanet, as expected, could only be partially overcome by excess UFH (**Fig. 1d**), achieving a level of anticoagulation less than what would be achieved with <1 U/mL UFH alone. This further validates that the ability of excess UFH to circumvent the effects of andexanet was dependent on the andexanet dose.

### Andexanet-associated heparin resistance increases thrombotic risk in extracorporeal circuits

To date, all laboratory descriptions of andexanet-associated heparin resistance have come from *in vitro* coagulation assays.^45^ However, *in vitro* coagulation tests do not necessarily reflect adequacy of anticoagulation or risk of excessive coagulation activation. Thus, to determine if andexanet actually prevents adequate anticoagulation for extracorporeal circulation, we studied the effects of andexanet on heparinization in a previously-described CPB model.^20, 39^ In this open-system model, human whole blood was continuously recirculated through a system that included a pump, venous reservoir, and membrane oxygenator. 4 U/mL UFH was added to blood prior to exposing it to the circuit. Following a one-hour circuit run, Andexanet was added to the circuit at varying concentrations and the circuit was run for an additional hour. Scanning electron microscopy (SEM) of the oxygenator membranes post circulation demonstrated no visible clot in the circuits anticoagulated with UFH alone (**Fig. 2a, Supplementary Fig. S1a**). However, all circuits to which andexanet were added showed extensive visible clot with SEM. Notably, as predicted from *in vitro* ACT and TEG studies, 16 U/mL UFH did not prevent visible clot formation in circuits supplemented with high dose andexanet. The addition of andexanet also resulted in overt compromise to flow in some of the extracorporeal circuits as evidenced by abrupt decreases in flow and increases in circuit pressure (**Fig. 2b-c, Supplementary Fig. S1b-c**), suggesting circuit obstruction by clot. Importantly, even circuits which remained patent had SEM evidence of clot formation.

**Figure 2.**
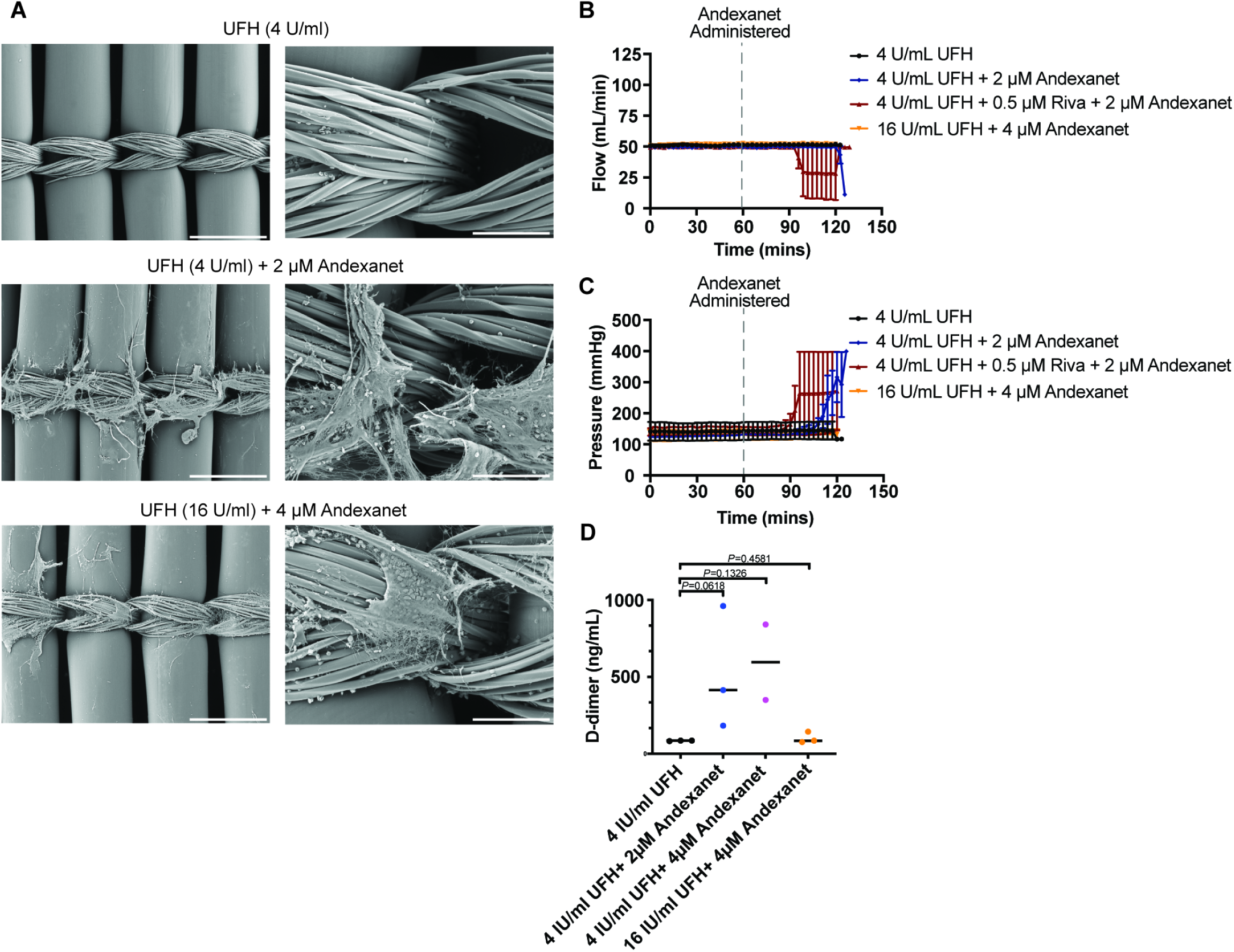
Andexanet-associated heparin resistance in extracorporeal circuits. (***a***) Representative SEM images (100x left and 500x right magnification, scale bars 500 μm left and 100 μm right) of oxygenator membranes following *ex vivo* circulation of blood heparinized with a typical UFH strategy and with UFH in addition to Andexanet. (***b,c***) Circuit flow (***b***) and pressure (***c***) in the *ex vivo* circuits under the indicated conditions for n=2-3 replicates (error bars shown represent 1 SD). (***d***) D-dimer levels measured in plasma obtained from *ex vivo* circuit blood following the experiment.

To evaluate how andexanet affects coagulation activation in UFH-anticoagulated circuits, we measured D-dimer levels following exposure to the CPB model. Compared to standard dose UFH-anticoagulated circuits, mean D-dimer levels were elevated in circuits exposed to standard dose UFH plus either dose of andexanet (**Fig. 2d**) although this approached but did not reach statistical significance. Surprisingly, despite SEM evidence of clot formation, D-dimer levels were not elevated in circuits anticoagulated with high dose UFH in the presence of high dose andexanet. The reason for this is unclear, but plasmin generation has been reported to be somewhat impaired by UFH.^46^ Those reports were performed with much lower concentrations of UFH than the 16 U/mL “high dose” UFH we used, and it is plausible that at such high doses, plasmin generation could be completely abrogated.

### Andexanet decreases AT inhibition of thrombin and FXa in the presence of UFH

Andexanet binds AT in the presence of heparin.^47^ We hypothesized that andexanet causes heparin resistance by preventing UFH-AT complexes from inhibiting thrombin or FXa. To test this, we measured the kinetics of UFH-AT inhibition of FXa and thrombin by measuring FXa-AT and thrombin-AT (TAT) complex formation. Andexanet dose-dependently slowed the rate of FXa inhibition by AT in the presence of 4 U/mL UFH (**Fig. 3a**). In the presence of 16 U/mL UFH (**Fig. 3b**), only high dose (4 μM) but not low dose (2 μM) andexanet impaired AT inhibition of FXa. Similarly, inhibition of thrombin by AT in the presence of heparin was slowed in the presence of andexanet (**Fig. 3c**).

**Figure 3.**
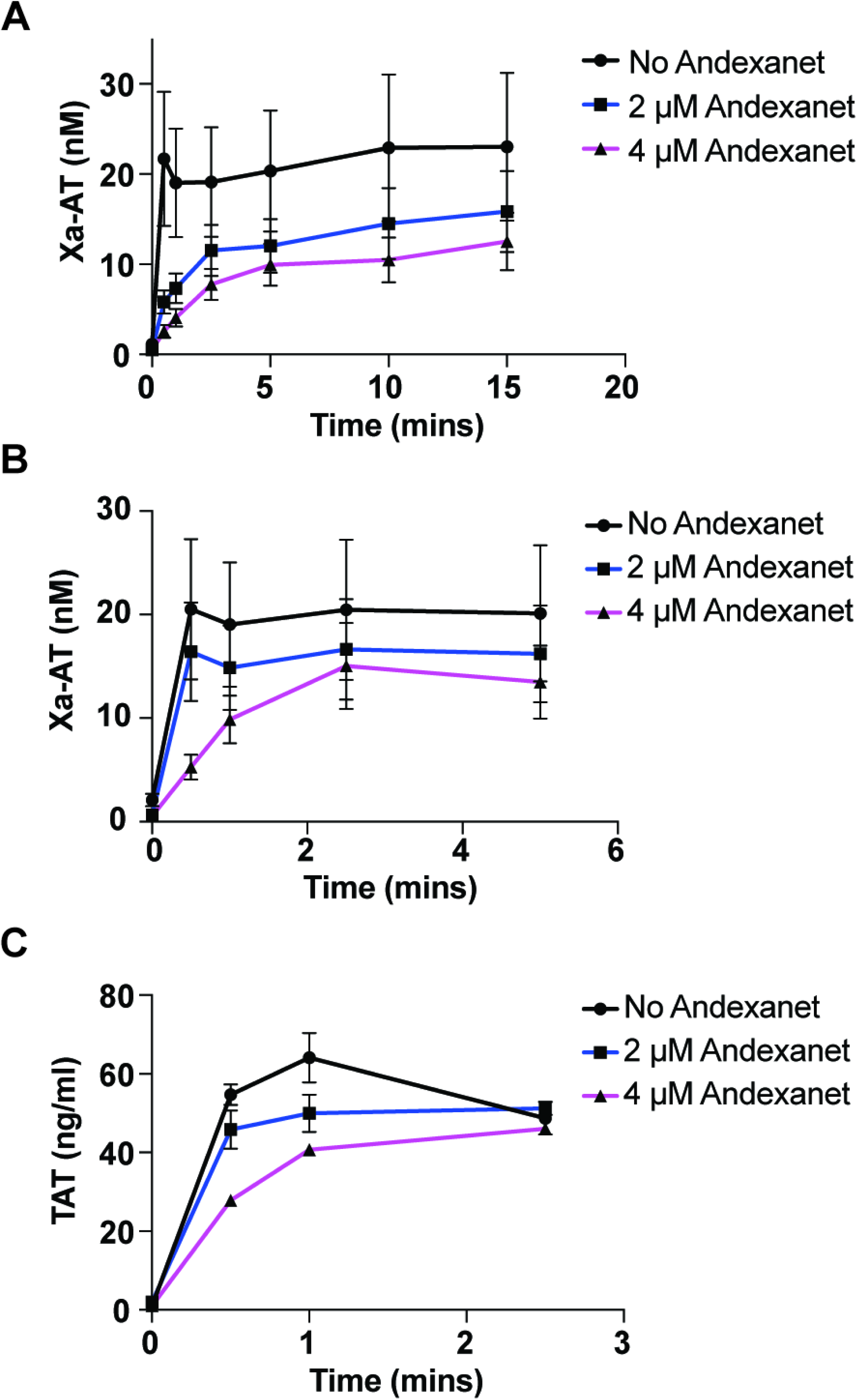
Effect of Andexanet on the kinetics of UFH-enhanced AT inhibition of FXa and thrombin. (***a,b***) Kinetics of FXa inhibition by AT in factor X-deficient human plasma in the presence of 4 U/mL (***a***) and 16 U/mL (***b***) UFH and the indicated concentrations of andexanet. (***c***) Kinetics of thrombin inhibition by AT in a purified system in the presence of 4 U/mL UFH and the indicated concentrations of andexanet. Experiments were performed in duplicate and results shown represent the mean ± SD.

While much of the effect of heparins on AT-mediated serine protease inhibition is due to direct effects of heparin on AT itself, other effects, including a bridging mechanism with respect to thrombin and, to a lesser extent, binding of heparin to the heparin binding exosite of FXa, have been reported.^48^ To determine if these other mechanisms of heparin-mediated serpin potentiation have a role in andexanet-associated heparin resistance, we repeated FXa-AT and TAT kinetics experiments with the minimal heparin pentasaccharide, fondaparinux (H5). Since H5 is too short to bind to both AT and the protease, if these other mechanisms were relevant, heparin resistance would be seen to a lesser extent (or not at all) with H5. As shown in **Supplementary Fig. S4**, the kinetics of H5-accelerated AT inhibition of FXa and thrombin were similar to those seen with UFH-accelerated AT inhibition, suggesting that the primary mechanism is due to conformational activation of the serpin itself rather than a template or bridging mechanism.

### Equilibrium modeling of the interaction between andexanet, UFH, and AT

While it is clear that andexanet-associated heparin resistance can cause subtherapeutic anticoagulation with UFH due to binding to UFH-AT complexes, it is not known what the role of different concentrations of each of these species plays. The expression in **Fig. 4a** summarizes the interaction between UFH, AT, and andexanet. The key species responsible for UFH-mediated anticoagulation is the free UFH-AT complex. When this complex binds to andexanet, it can no longer exert its anticoagulant function. We hypothesized that the relative concentrations of andexanet, UFH, and AT are the primary drivers of free UFH-AT levels and therefore the degree of anticoagulation. To test this hypothesis, we used the previously described rate constants of these interactions to model andexanet binding to UFH-AT complexes.^36, 37^ These “heparin response curves” provide a graphical representation of the impact of andexanet, UFH, and AT concentrations on the equilibrium concentration of UFH-AT complexes.

**Figure 4.**
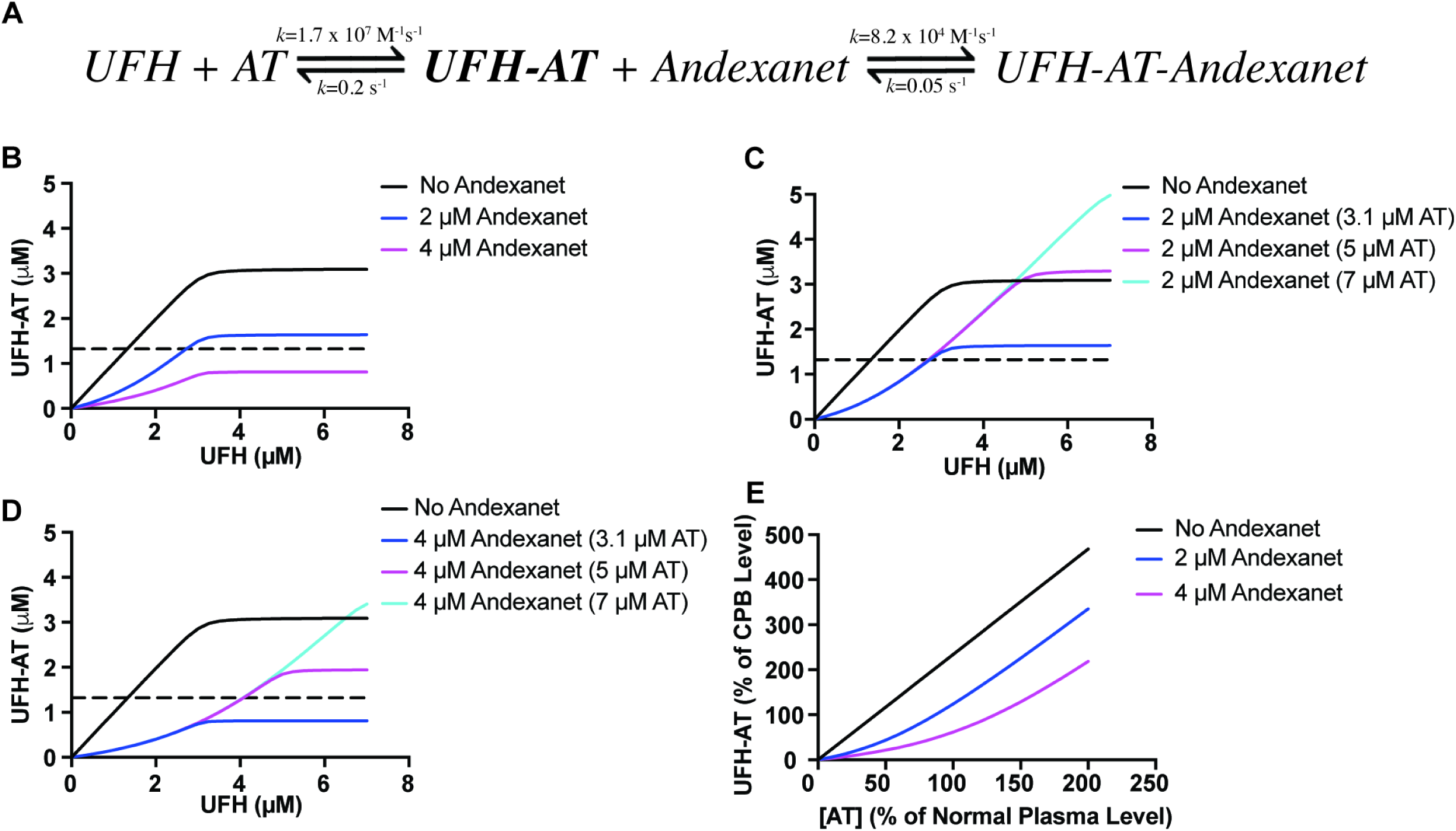
Equilibrium modeling of the effects of Andexanet on UFH binding to AT. (***a***) Equilibrium expression for UFH binding to AT and andexanet binding to the UFH-AT complex. The previously determined association and dissociation constants for each expression are shown. (***b***) Effect of UFH on the steady-state concentration of the UFH-AT complex at the concentrations of andexanet shown, assuming a starting concentration of 3.1 μM AT (normal plasma AT concentration). (***c,d***) Effect of UFH on the steady-state concentration of the UFH-AT complex at normal and supraphysiologic AT levels in the presence of (***c***) 2 μM and (***d***) 4 μM andexanet. (***e***) Effect of plasma AT concentration on the maximum (assuming infinite UFH levels) achievable steady-state UFH-AT complex levels, expressed as a percentage of the level of UFH-AT complex needed for safe initiation of CPB. The dashed lines in panels ***b-d*** represent the UFH-AT concentration produced by normal CPB concentrations of UFH.

In the absence of andexanet, heparin response is biphasic: nearly linear UFH-AT complex formation with increasing concentrations of UFH until AT is exhausted (**Fig. 4b, black line**). In the presence of 2 μM andexanet, heparin response is triphasic: 1) minimal free UFH-AT formation when UFH concentration is below andexanet concentration, 2) nearly normal heparin responsiveness with a similar slope to the native heparin response curve when the UFH concentration is greater than the andexanet concentration, and 3) exhaustion of AT with no additional UFH-AT formation (**Fig. 4b, blue line**). Notably, the maximal free UFH-AT concentration achieved (“ceiling”) is lower due to the rightward shift of the curve and limitation by AT. However, this ceiling is approximately the expected level of UFH-AT complex needed for therapeutic anticoagulation for CPB (**Fig. 4b, dashed line**), which may explain why heparin resistance due to low dose andexanet can be overcome with additional UFH. In the presence of 4 μM andexanet, only two phases are seen: since andexanet concentration exceeds AT concentration, when andexanet is exhausted no residual AT is available so the middle phase of robust heparin responsiveness does not occur (**Fig. 4c, magenta line**). This results in a ceiling well below the level needed for CPB.

To evaluate the potential impact of AT supplementation on andexanet-associated heparin resistance, we modeled heparin responsiveness at various plasma AT concentrations. In the presence of 2 μM andexanet, supraphysiologic AT concentrations simply extended the second phase of the triphasic equilibrium as more UFH was needed to exhaust the higher AT levels (**Fig. 4c**). This suggests that supplementation of AT in the presence of 2 μM andexanet is unlikely to be of much benefit to achieve therapeutic levels of anticoagulation for CPB. However, in the presence of 4 μM andexanet, supraphysiologic AT concentrations restore the triphasic equilibrium not seen at physiologic AT levels (**Fig. 4d**). This suggests that supplementation of AT (in addition to high dose UFH) is necessary at high andexanet concentrations to enable heparin responsiveness, as it allows the AT concentration to exceed the andexanet concentration.

Equilibrium modeling indicates that the relative concentrations of AT and andexanet are critical determinants of heparin responsiveness. Patients presenting for emergent cardiac surgery often have decreased levels of AT.^49^ To determine the impact of AT level on ceiling heparin responsiveness, we modeled the ceiling heparin response with respect to plasma AT level (**Fig. 4e**). Andexanet decreased the ceiling heparin response at all AT levels. Importantly, the ceiling heparin response in the presence of 2 μM andexanet is only sufficient for CPB at or above 90% normal AT levels. Even at normal AT concentrations, the ceiling heparin effect is inadequate when 4 μM andexanet is present and is only adequate for CPB when AT levels are at least 140% of normal levels.

### Validation of equilibrium model using thrombin generation studies

Equilibrium modeling of andexanet-associated heparin resistance reveals a triphasic heparin response if the andexanet concentration does not exceed the AT concentration. To validate this triphasic equilibrium, we measured thrombin generation in normal human plasma supplemented with andexanet and UFH. Heparin responsiveness was robust in the absence of andexanet (**Fig. 5a, Supplementary Fig. S5**). No plateau (AT depletion) phase was seen, likely due to the assay’s high sensitivity for UFH. In the presence of 2 μM andexanet, we observed the predicted triphasic response to increasing UFH concentration with a robust phase of heparin responsiveness between 8 and 12 U/mL UFH (approximately 2.7 to 4 μM assuming 200 U/mg UFH). Thrombin generation in the presence of 4 μM andexanet only displayed the first and third phases, consistent with our prediction that if andexanet concentration exceeds AT concentration, the normal second phase will not occur.

**Figure 5.**
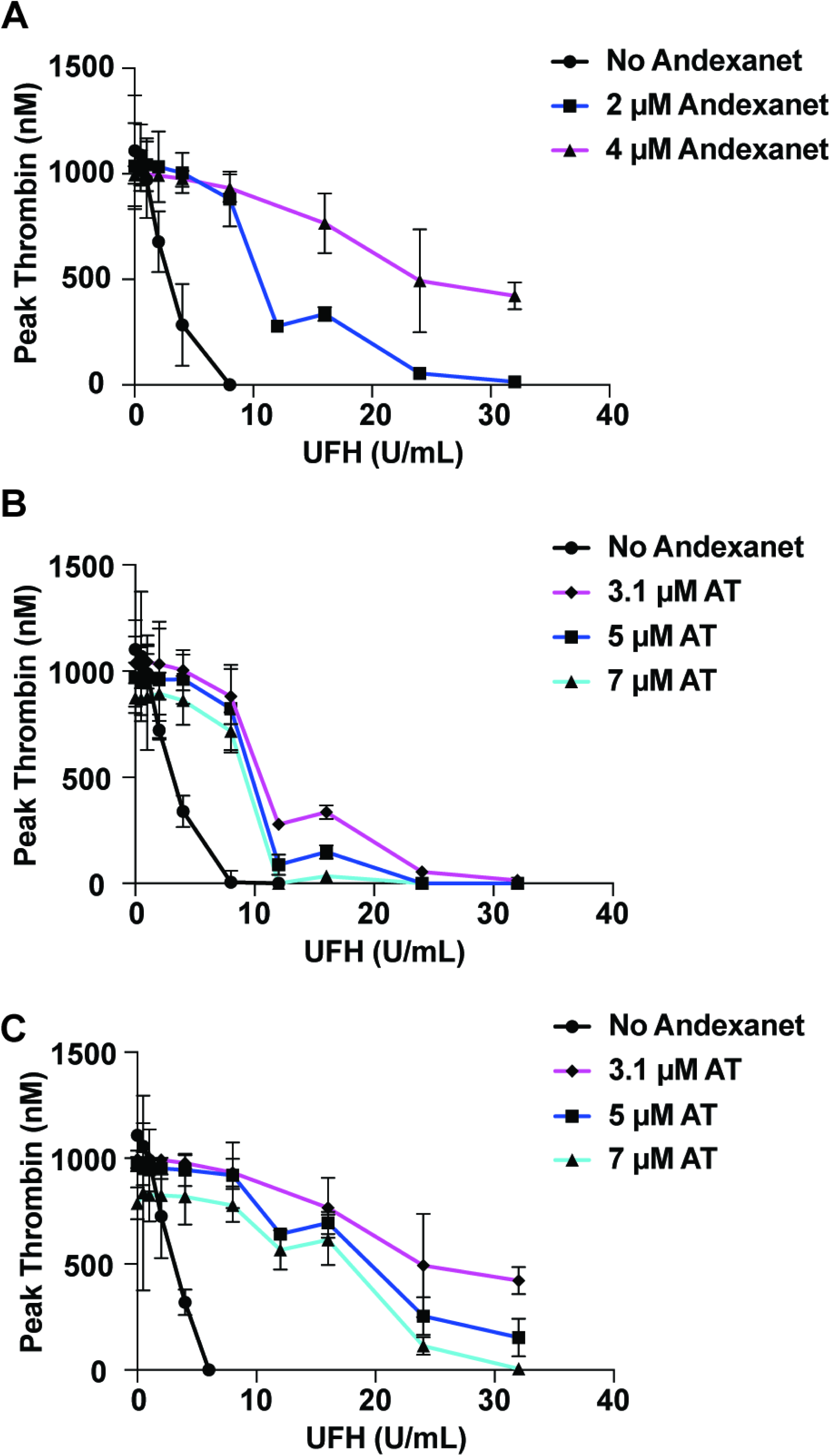
Effect of Andexanet on UFH inhibition of thrombin generation in human plasma. ***(a)*** Inhibition of thrombin generation by UFH in normal human plasma and in the presence of the indicated concentrations of andexanet. (***b,c***) Inhibition of thrombin generation by UFH in normal human plasma (3.1 μM AT) and at the indicated supraphysiologic concentrations of AT in the presence of (***b***) 2 μM and (***c***) 4 μM andexanet. In all experiments, results are plotted as the mean ± SEM of experiments performed in duplicate. Raw thrombin generation curves are shown in **Supplementary Figures S5-S7.**

This model also predicts that when andexanet concentration is less than the AT concentration, supplementing AT will not affect the starting UFH concentration of the second phase, but will affect ceiling heparin effect, since there is additional excess AT. When additional AT was added to normal human plasma containing 2 μM andexanet, the expected triphasic heparin response was again seen, with little-to-no change in beginning of the second phase (**Fig. 5b, Supplementary Fig. S6**). However, the heparin effect at the end of the second phase was greater in the AT-supplemented experiments.

As described, the second phase was absent in the presence of 4 μM andexanet due to the fact that the andexanet concentration exceeded the AT concentration. When AT was supplemented to increase AT concentration above that of andexanet, a second phase appeared, albeit at approximately twice the UFH concentration (16 to 24 U/mL UFH) as seen in the presence of 2 μM andexanet (**Fig. 5c, Supplementary Fig. S7**). This is consistent with our model, which predicts that AT supplementation will restore heparin responsiveness at very high UFH concentrations.

## Discussion

The ability to rapidly reverse the effects of an anticoagulant drug is desirable, as it increases the versatility and safety of the anticoagulant. Most reversal agents for direct oral anticoagulants, including andexanet, are drug-sequestering antidotes.^11, 50^ In order to accommodate the wider chemical diversity of direct FXa inhibitors, andexanet was created as a variant of FXa itself.^11^ This is an attractive feature of andexanet, since it is a single drug that can reverse the effects of all commercially available direct FXa inhibitors. However, the approach of using a nonfunctional coagulation factor variant increases the risk of off-target effects compared to other strategies (such as antibody fragments) because the molecule retains many of its binding properties. Two properties of andexanet increase the probability of unintended or off-target effects. First, in the active site, andexanet is identical to that of FXa with the exception of a single amino acid change to prevent amidolytic activity.^11^ This allows andexanet to retain many of FXa’s active site properties (and therefore bind direct FXa inhibitors). However, the active site of FXa is an important binding site for regulatory plasma protease inhibitors including tissue factor pathway inhibitor (TFPI) and AT and, indeed, Andexanet has been shown to deplete TFPI.^16, 51^ Depletion of TFPI, which is present in plasma at 0.1% the concentration of low dose andexanet,^52^ has been used to explain, in part, why the effects of andexanet on restoring thrombin generation seem to persist longer than the drug’s effect on anti-FXa activity.^16^ While this may be of benefit to the reversal agent properties of andexanet, TFPI also plays a crucial role in prevention of disseminated intravascular coagulation, and its depletion could lead to thrombotic complications, especially in cardiac surgery. The second factor contributing to potential off-target effects of andexanet is the high concentration of andexanet needed for reversal. Because therapeutic plasma concentrations of direct FXa inhibitors are in the high nanomolar range, andexanet is administered to achieve micromolar concentrations to ensure adequate anticoagulant reversal.^16^ At these concentrations, andexanet levels approach or, in fact, exceed those of many coagulation components, including AT and TFPI.

Although early pre-clinical data indicated that andexanet does not meaningfully bind to AT in the absence of heparin, it was shown to bind to AT when heparin is present.^47^ It was suggested that this would enable andexanet to also reverse indirect FXa inhibitors such as UFH, fondaparinux, and low molecular weight heparins.^11^ The observation that andexanet binds AT only in the presence of heparin is a reflection of the well-established two-step mechanism by which AT inhibits its target serine proteases. In the first step, the “reactive center loop (RCL)” of AT binds the active site of its target to form a non-covalent Michaelis complex. The protease then attempts to cleave the RCL which results in a covalent complex between AT and its now irreversibly-denatured target.^53^ The affinity of AT for FXa and thrombin in the noncovalent first step is incredibly low, likely in the millimolar range.^21^ Thus, the reaction only occurs because the irreversible, covalent second step drives it to completion. Heparins make the otherwise weak first Michaelis binding step much tighter, thus accelerating AT inhibition of its targets. Because andexanet lacks the catalytic S195 needed to attack the RCL and covalently bind to AT, it can only interact with AT via the very weak Michaelis complex. Since heparin enhances the noncovalent affinity of AT for its targets, andexanet (in the presence of heparin), despite lacking S195, binds AT with high nanomolar affinity (130-710 nM).^37, 47^ This explains why andexanet only binds AT in the presence of heparin.

Multiple etiologies of heparin resistance have been described including nonspecific heparin binding and AT deficiency. Our data reveal that andexanet-associated heparin resistance has distinct properties compared to traditional forms of heparin resistance. Unlike in traditional heparin resistance, increasing heparin doses further depletes AT by allowing more AT binding to andexanet. Additionally, as shown in **Fig. 4c-d**, supplementing AT at typical UFH concentrations does little to improve heparin responsiveness. Instead, high doses of UFH must be used to first consume all andexanet present (at the expense of AT depletion). If any AT remains after andexanet is consumed, additional UFH will potentiate it, but if the residual amount of AT is inadequate, AT must be supplemented as well.

Patients requiring emergent cardiac surgery often cannot wait for adequate anticoagulant washout. While patients on warfarin can have their anticoagulation reversed prior to emergent cardiac surgery, ideally patients receiving direct factor Xa inhibitors should not receive andexanet for anticoagulant reversal prior to CPB as this will result in heparin resistance. Unfortunately, pre-CPB andexanet administration is unlikely to be completely avoidable, especially as the most recent societal guidelines recommend its administration.^32^ Patients who develop pericardial effusions following electrophysiologic procedures may be given andexanet to prevent further expansion of the pericardial effusion and subsequent pericardial tamponade, but this will complicate heparinization should CPB become necessary. Additionally, providers less familiar with andexanet-associated heparin resistance may administer andexanet to patients who need emergent CPB. Finally, following CPB, a subset of patients will require urgent or emergent return to CPB for technical or hemodynamic reasons. If a patient receives andexanet following CPB and then needs to emergently return to CPB, this could be particularly problematic as empiric dosing of heparin may be insufficient due to andexanet-associated heparin resistance. In all of these scenarios, one option is to use a parenteral direct thrombin inhibitor such as bivalirudin or argatroban to achieve anticoagulation for CPB. However, whereas UFH can be reversed with protamine sulfate, these agents lack a reversal agent. Thus while they have been used safely for CPB, the inability to rapidly eliminate the anticoagulant effect of direct thrombin inhibitors following CPB may dissuade some providers from their use.^54^

For these reasons, it is important to understand the variables that determine heparin responsiveness following andexanet administration. Equilibrium modeling of the expression shown in **Fig. 4a** results in a triphasic model that illustrates the importance of the relative concentrations of andexanet and AT. Specifically, it suggests that patients with andexanet-associated heparin resistance may fall into two distinct groups: those whose andexanet levels are below the AT concentration, and those whose andexanet levels are above the AT concentration. In general, patients in the former group may have some degree of heparin responsiveness in the “second phase” of the heparin response curve, whereas those in the latter group would be expected to lack significant heparin responsiveness. The fact that patients receiving andexanet could fall into two different patterns of heparin responsiveness may help to explain why some case reports described achieving therapeutic heparinization with heparin alone while others reported requiring AT supplementation.^27–31^ It would be beneficial to know which group a patient falls into as this could help predict what dose of UFH might be needed and whether AT supplementation is also required. However, determining this is not practical since andexanet dosing is not adjusted for weight and volume of distribution may vary, which will affect andexanet concentration. Moreover, patients presenting for emergent cardiac surgery have lower AT levels than those undergoing elective cardiac surgery.^49^ As shown in **Fig. 4e**, even if plasma andexanet concentration could be predicted precisely, a patient with only slight AT deficiency (below 90% of normal levels) would not be expected to achieve therapeutic anticoagulation with high dose UFH alone. This work also demonstrates why andexanet-associated heparin resistance is not limited to high-anticoagulation applications. Very little heparin response will occur during the first phase of the equilibrium, and thus the amount of UFH needed to overcome the effects of andexanet will not scale proportionately to the level of anticoagulation desired.

Because of the *in vitro* nature of this work, we cannot make clinical recommendations regarding patient management in the event that andexanet-associated heparin resistance is encountered. However, our findings suggest several specific clinical questions that warrant further investigation. First, in a patient who has recently received andexanet, how much UFH should be administered before consideration of AT administration? The 500 U/kg UFH cutoff commonly used for AT-mediated heparin resistance may warrant reevaluation in the context of andexanet since, as shown in **Fig. 4c** and **Fig. 5b**, additional AT without sufficient UFH may not significantly increase the level of anticoagulation. A second, related question is, in the event that AT supplementation is needed, what dose should be used? Again, our work suggests that this may be different from heparin resistance due to AT-deficiency and may be highly variable depending on the exact concentrations of andexanet and AT in the patient’s plasma. Finally, this work suggests that there may be a role for the development of a rapid, specific assay to assess plasma andexanet concentrations so that it can be compared to AT levels.

The pharmacokinetics and pharmacodynamics of andexanet with respect to andexanet-associated heparin resistance also warrant further evaluation. Andexanet reverses the effects of apixaban and rivaroxaban with an approximately 1-hour pharmacodynamic half-life.^15^ This could suggest that andexanet-associated heparin resistance might be short-lived. However, pharmacokinetic studies of andexanet demonstrate that andexanet has a half-life of 4.35-7.5 hours, far longer than its pharmacodynamic half-life.^55^ It is not entirely clear which half-life is more relevant to andexanet-associated heparin resistance, and further investigation is necessary to determine how long the problem of andexanet-associated heparin resistance persists after administration.

### Conclusions

Andexanet-associated heparin resistance is a major impediment to the use of andexanet in the periprocedural period. Despite increasing awareness of this phenomenon among clinicians, andexanet administration in patients who may require heparinization will occasionally occur. By elucidating the mechanism, this report provides insight into the numerous case reports of andexanet-associated heparin resistance and creates a biochemical foundation that can guide further clinical investigation of the issue.

## Acknowledgements

## Acknowledgements

We thank Drs. Rodney Camire, Lindsey George, and Sriram Krishnaswamy for their critical appraisal of the manuscript and useful suggestions.

## Supplemental Material

Detailed Methods

Table S1

Figures S1-S7

Supplemental References

Major Resources Table

